# *C9orf72*-associated arginine-rich dipeptide repeats induce RNA-dependent accumulation of Staufen in nucleus

**DOI:** 10.1101/773796

**Authors:** Eun Seon Kim, Chang Geon Chung, Yoon Ha Kim, In Jun Cha, Jeong Hyang Park, Jaekwang Kim, Chang Man Ha, Hyung-Jun Kim, Sung Bae Lee

## Abstract

Accumulation of RNA in the nucleus is one of the pathological features of *C9orf72-*associated amyotrophic lateral sclerosis and frontotemporal dementia (C9-ALS/FTD), yet its potential toxic cellular consequences remain largely undefined. RNA accumulated in the nucleus may interact with and increase nuclear localization of RNA-binding proteins (RBPs). Here, we show in C9-ALS/FTD *Drosophila melanogaster* model that Staufen, a double-stranded RBP normally localized in cytoplasm, accumulates in the nucleus, which is in contrast to many nuclear-localized RBPs, such as TDP-43 and FUS, whose cytoplasmic accumulation is thought to be a pathological hallmark of ALS/FTD. We found that in *Drosophila* neurons expressing arginine-rich dipeptide repeat proteins (DPRs), Staufen accumulated in the nucleus in an RNA-dependent manner. In the nucleus, Staufen localized closely to, and potentially interacts with, heterochromatin and nucleolus in *Drosophila* C4 da neurons expressing poly(PR), a proline-arginine (PR) DPR. PR toxicity in C4 da neurons increased Fibrillarin staining in the nucleolus, which was enhanced by *stau* heterozygous mutation. Furthermore, knockdown of *fib* exacerbated retinal degeneration mediated by PR toxicity, which suggests that increased amount of Fibrillarin by *stau* heterozygous mutation is protective. Heterozygotic mutation of *stau* could also mitigate retinal degeneration and rescue viability of flies exhibiting PR toxicity. Taken together, our data show that nuclear accumulation of cytoplasmic protein, such as Staufen, may also be an important pathological feature of C9-ALS/FTD.

**Author summary:** Cytoplasmic accumulation of nuclear RNA-binding proteins (RBPs) is one of the common pathological features of amyotrophic lateral sclerosis (ALS) and frontotemporal dementia (FTD). In *C9orf72*-associated ALS/FTD fly model, we found that Staufen, a double-stranded (ds) RBP normally localized mostly in cytoplasm, accumulates in the nucleus in an RNA-dependent manner. Next, we checked wherein the nucleus Staufen accumulates and found that Staufen partially co-localizes with heterochromatin and nucleolus. Interestingly, the expression of Fibrillarin, a nucleolar protein, was significantly increased by *C9orf72*-derived PR toxicity and further augmented by reduction in *stau* dosage, the gene encoding Staufen. When we knocked down *fib*, the gene encoding Fibrillarin, PR-induced retinal degeneration was exacerbated. This indicates that increased Fibrillarin expression by *stau* dosage reduction is protective. Furthermore, when we reduced *stau* dosage in flies presenting PR toxicity, their retinal degeneration and viability were largely rescued. Based on these data, we suggest that nuclear accumulation of Staufen is an important feature of C9-ALS/FTD and suggest that reducing *stau* dosage is a promising therapeutic target.

## Introduction

Amyotrophic lateral sclerosis (ALS) and frontotemporal dementia (FTD) are two diseases on a single pathogenic spectrum with overlapping genetic etiologies [1]. GGGGCC (G_4_C_2_) repeat expansion mutation in the intron of *C9orf72* is the most common cause of both familial and sporadic cases of ALS and FTD (C9-ALS/FTD) [2, 3]. G_4_C_2_ repeat expansion produces both sense (G_4_C_2_) and antisense (G_2_C_4_) RNAs that can either form toxic nuclear foci [4, 5] or undergo repeat-associated non-ATG translation (RANT) in the cytoplasm to generate five types of dipeptide repeat proteins (DPRs) [6, 7]: glycine-alanine (GA), glycine-proline (GP), glycine-arginine (GR), proline-arginine (PR), and proline-alanine (PA). Among them, arginine-rich DPRs have been shown to be most toxic in fly and mammalian models for ALS [8].

In recent years, nucleocytoplasmic transport defect, which includes decreases in protein import into and mRNA export from the nucleus, has been identified as one of the prominent pathogenic features of C9-ALS/FTD [9–12], leading to accumulation of proteins in the cytoplasm and mRNA in the nucleus, respectively. In addition, recent studies showed nuclear accumulation of double-stranded RNA (dsRNA) in *tdp-1*-deleted *C. elegans* [13] and in PR-expressing mouse cortical neurons [14]. Currently, potentially toxic consequences of decreased protein import are actively being explored, one of which is cytoplasmic mis-localization of TDP-43 [15], a pathological hallmark of ALS/FTD [16]. However, potentially toxic consequences of RNA accumulation in the nucleus remain largely underexplored.

RNA-binding proteins (RBPs) such as hnRNPA1 have been shown to egress out of the nucleus to the cytoplasm when there is a decreased level of nuclear RNA [17, 18]. These studies lead to a hypothesis that increased RNA levels in the nucleus might induce nuclear retention of certain RBPs, resulting in their subsequent gain of nuclear toxicity in ALS. Thus, to test our hypothesis, we used C9-ALS/FTD *Drosophila* model system to screen for and identify RBPs whose nuclear accumulation leads to neurotoxicity and to understand its underlying mechanisms.

## Results

### Arginine-rich DPR proteins increase nuclear accumulation of Staufen in neurons

To identify potential RNA-binding proteins (RBPs) that show increased nuclear localization in C9-ALS/FTD *Drosophila* model, we screened in C4 da neurons, which was recently used to model ALS in flies [19], for RBPs whose accumulation in the nucleus was increased compared to the controls. For the screen, we used as C9-ALS/FTD *Drosophila* model the flies that expressed PR repeat proteins (V5-PR36) and not GR repeat proteins, because recent studies showed that between the two arginine-rich DPRs, PR is more closely associated with nucleocytoplasmic transport defects [10, 20, 21]. In addition, by using V5-PR36, which is expressed from an alternative codon different from repeated G_4_C_2_ sequence, we were able to preclude detection of RBPs mis-localized by G_4_C_2_ RNA from the screen.

The genetic screen identified Staufen as the RBP whose accumulation in the nucleus increased most significantly. In control (CTRL) C4 da neurons, GFP-tagged Staufen showed mostly cytoplasmic localization with small puncta in the nucleus, whereas in C4 da neurons expressing V5-PR36, GFP-Staufen puncta were much more prominent in the nucleus (Fig 1A). We then quantitatively measured the mean intensity (y-axis) and counted pixel numbers (x-axis) for nuclear GFP-Staufen puncta in controls and in C4 da neurons expressing V5-PR36 and plotted their relative values (Fig 1B). V5-PR36-expressing C4 da neurons showed puncta with substantially higher mean fluorescence intensity (i.e. higher density) and pixel number (i.e. larger size) compared to those in controls. To estimate relative amount of GFP-Staufen proteins in the nucleus of neurons with or without V5-PR36 expression, we multiplied the mean fluorescence intensity with the pixel number of GFP-Staufen puncta. C4 da neurons expressing V5-PR36 showed significantly (p = 0.0022) higher amount of Staufen proteins in the nucleus compared to the controls (Fig 1C).

**Fig 1.**
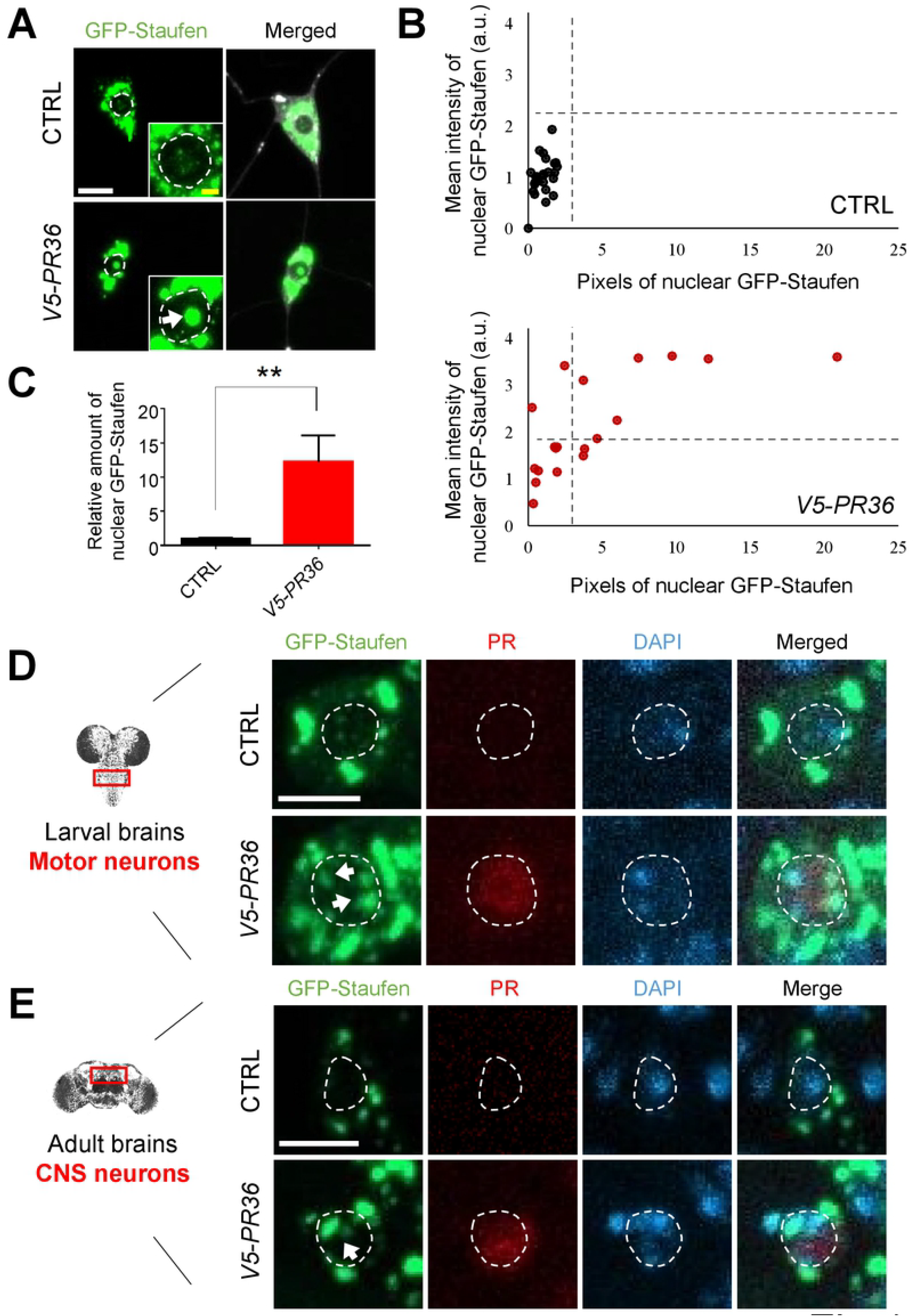
PR toxicity increases nuclear accumulation of Staufen in neurons. (A) V5-PR36 expression in C4 da neurons of third instar larvae significantly increased nuclear accumulation of GFP-Staufen (Genotypes. CTRL: *UAS-GFP-stau/+*; *PPK*^*1a*^-*gal4>UAS-mCD8RFP/+*, *V5-PR36*: *UAS-GFP-stau/+*; *PPK*^*1a*^-*gal4>UAS-mCD8RFP/UAS-V5-PR36*). Bottom right insets are magnified images of the nuclei. Dashed circular lines (white) outline the nuclei. Arrow (white) indicates increased nuclear GFP-Staufen puncta. Scale bars (yellow), 10 µm; (white), 2 µm. (B) Individual nuclear GFP-Staufen puncta in C4 da neurons with denoted genotypes from Fig. 1a were counted and plotted. y-axis: relative mean intensity of GFP-Staufen puncta in nucleus; x-axis: relative pixel number of GFP-Staufen puncta in nuclei. n≥19 GFP-Staufen puncta (C) The pixel number and the mean intensity of nuclear GFP-Staufen puncta (obtained from Fig 1B) were multiplied to provide an estimate of relative amount of nuclear GFP-Staufen. **p = 0.0022 by two-tailed student’s t-test; error bars, ± SEM; n≥19 GFP-Staufen puncta. (D) IHC of dissected third instar larval motor neurons shows that V5-PR36 expression increased nuclear accumulation of GFP-Staufen (Genotypes. CTRL: *UAS-GFP-stau/+*; *D42-Gal4>UAS-mCD8RFP/+*, *V5-PR36*: *UAS-GFP-stau*/*+*; *D42-Gal4>UAS-mCD8RFP*/*UAS-V5-PR36*). Dashed circular lines (white) outline the nuclei. Arrows (white) indicate stronger nuclear GFP-Staufen signal compared to the control. Scale bar (white), 5 µm. (E) IHC of adult central nervous system (CNS) neurons expressing V5-PR36 shows an increase in nuclear accumulation of GFP-Staufen (Genotypes. CTRL: *UAS-GFP-stau/+*; *elavGS-gal4/+*, V5-*PR36*: *UAS-GFP-stau/+*; *elavGS-gal4/UAS-V5-PR36*). The transgenic expression was induced by feeding 100 µM of RU486 for 20 days after eclosion at 27 ℃. Dashed circular lines (white) outline the nuclei. Arrow (white) indicates stronger nuclear GFP-Staufen signal compared to the controls. Scale bar (white), 5 µm.

To investigate whether nuclear accumulation of Staufen is specific to V5-PR36 expression, we examined in C4 da neurons the localization pattern of GFP-Staufen after expressing other C9-ALS/FTD constructs: G_4_C_2_36, G_4_C_2_36-RNA only, Myc-PA36, and HA-GR36. Among them, G_4_C_2_36 and HA-GR36 expression led to increased nuclear accumulation of Staufen (S1A-C Fig), whereas G_4_C_2_36-RNA only and Myc-PA36 expression did not. These results suggest that only the constructs that produce arginine-rich DPRs (i.e. G_4_C_2_36, V5-PR36, and HA-GR36), but not the others (i.e. G_4_C_2_36-RNA only and Myc-PA36), induce nuclear accumulation of Staufen. Next, we questioned whether V5-PR36 expression can increase nuclear accumulation of Staufen in other neuron types. To this end, we expressed V5-PR36 in motor neurons as well as in the entire central nervous system using *D42-gal4* and *elavGS*-*gal4* drivers, respectively. Consistent with our results from C4 da neurons, expression of V5-PR36 in other neuron types in *Drosophila* brains showed an increase in nuclear accumulation of Staufen (Fig 1D and 1E), compared to controls. These data collectively suggest that arginine-rich DPRs induce nuclear accumulation of Staufen in *Drosophila* neurons.

### Nuclear-accumulated Staufen partially co-localizes with heterochromatin in PR36-expressing neurons

A recent study reported that PR repeat proteins co-localize with heterochromatin and disrupt heterochromatin protein 1α (HP1α) liquid droplets [14]. This disruption in mouse cortical neurons expressing PR repeat proteins leads to mis-expression of repetitive elements (RE) that form dsRNA, which accumulates within the nucleus [14]. We hypothesized based on this previous study that Staufen, which is a dsRNA-binding protein (dsRBP), may be recruited to the heterochromatin region, a site of dsRNA production, in neurons presenting PR36 toxicity. To test this hypothesis, we expressed RFP-tagged HP1, a *Drosophila* ortholog of HP1α, and compared its co-localization pattern with GFP-Staufen with or without PR36 expression in C4 da neurons (Fig 2A and 2B). C4 da neurons with PR36 expression showed that greater fraction of HP1 puncta co-localized with GFP-Staufen puncta than in controls (p = 0.0339) (Fig 2C). Consistently, GFP-Staufen partially co-localized with 4′, 6-diamidino-2-phenylindole (DAPI), which labels heterochromatin, in C4 da neurons expressing V5-PR36 (Fig 2D and E). These data suggest that PR toxicity can augment heterochromatin distribution of Staufen in *Drosophila* neurons.

**Fig. 2.**
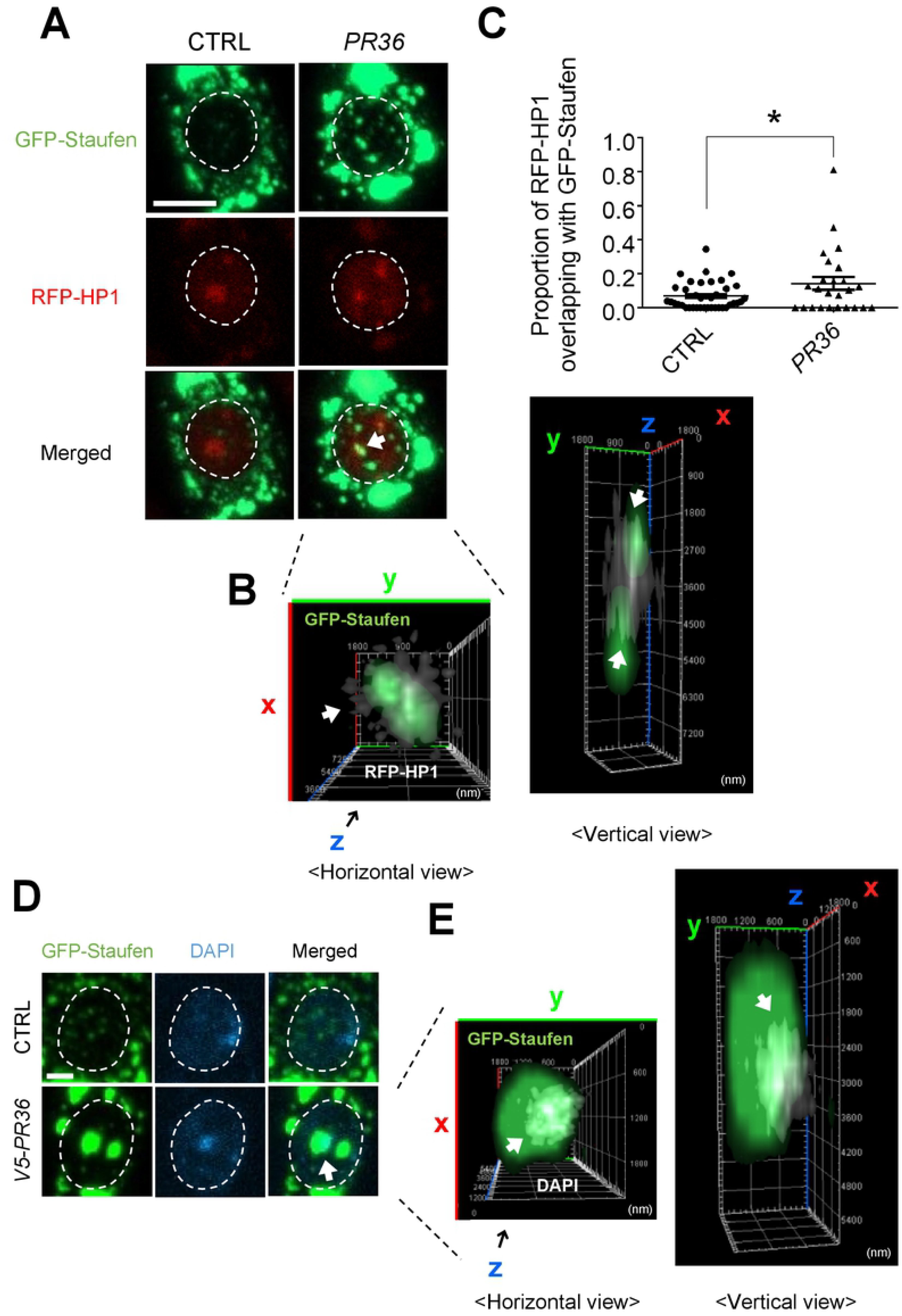
Nuclear-accumulated Staufen partially co-localizes with heterochromatin in C4 da neurons expressing PR36. (A) Nuclear-accumulated GFP-Staufen partially co-localizes with RFP-HP1 in V5-PR36-expressing C4 da neurons (Genotypes. CTRL: *UAS-GFP-stau/+; PPK*^*1a*^-*gal4/RFP-HP1*, *PR36*: *UAS-GFP-stau/UAS-PR36; PPK*^*1a*^-*gal4/RFP-HP1*). Dashed circular lines (white) outline the nuclei. Arrow (white) indicates co-localization between nuclear GFP-Staufen and RFP-HP1. Scale bar, 5 µm. (B) 3D images of Fig 2B are presented to better visualize the overlap of GFP-Staufen and RFP-HP1 in the nucleus. Arrows (white) indicate co-localization between nuclear GFP-Staufen and RFP-HP1 (white). (C) The proportion of RFP-HP1 puncta that co-localize with GFP-Staufen were individually measured and plotted. *p = 0.0339 by two-tailed student’s t-test; error bars, ± SEM; n≥25 RFP-HP1 puncta. (D) IHC of dissected third instar larvae shows that nuclear-accumulated GFP-Staufen partially co-localizes with DAPI in V5-PR36-expressing C4 da neurons (Genotypes. CTRL: *UAS-GFP-stau/+; PPK*^*1a*^-*gal4>UAS-mCD8RFP/+*, *V5-PR36*: *UAS-GFP-stau/+; PPK*^*1a*^-*gal4>UAS-mCD8RFP/UAS-V5-PR36*). DAPI (blue) staining labels heterochromatin. Dashed circular lines (white) outline the nuclei. Arrow (white) indicates co-localization between nuclear GFP-Staufen and DAPI. Scale bar, 2 µm. (E) 3D images of Fig 2D are presented to better visualize the overlap of GFP-Staufen and DAPI in the nucleus. Arrows (white) indicate co-localization between nuclear GFP-Staufen and DAPI (white).

Next, we asked how PR toxicity might induce nuclear accumulation of dsRNA in C4 da neurons. Recent studies showed that loss of nuclear TDP-43, a prominent pathological feature of C9-ALS/FTD [22], can derepress RE transcription from heterochromatin [13, 23, 24], leading to accumulation of dsRNA in the nucleus. Thus we tested whether decreasing the amount of nuclear TBPH, a *Drosophila* ortholog of TDP-43, is sufficient to increase accumulation of Staufen in the nucleus. To this end, we knocked down *TBPH* and examined the localization pattern of Staufen in the nucleus of C4 da neurons (S2A Fig). Compared to the controls, *TBPH* RNAi led to increased accumulation of Staufen in the nucleus of C4 da neurons (S2B-D Fig). These data suggest that PR toxicity might increase nuclear accumulation of Staufen through reducing nuclear TBPH.

### Increased nuclear accumulation of Staufen by PR toxicity is RNA-dependent

It is possible that Staufen can accumulate in the nucleus by its binding to RNAs, such as dsRNA. To test whether PR-induced nuclear accumulation of Staufen is dependent on RNA, we dissected larvae with V5-PR36 expression in C4 da neurons and treated them with RNase A for 1hr at 37 ℃ before fixing them for imaging analysis. The RNase A treatment protocol we used effectively reduced RNA levels in C4 da neurons (S3A and S3B Fig). Reducing RNA levels led to a substantial decrease in the V5-PR36-induced nuclear accumulation of Staufen (Fig 3A). We then measured the mean intensity (y-axis) and counted pixel numbers (x-axis) of nuclear GFP-Staufen puncta in C4 da neurons expressing V5-PR36 with or without RNase A treatment and plotted their relative values (Fig 3B). RNase A treatment led to decreases in density and size of GFP-Staufen puncta. Relative amount of GFP-Staufen proteins in the nucleus was calculated by multiplying the mean fluorescence intensity with the pixel number of GFP-Staufen puncta. RNase A-treated C4 da neurons expressing V5-PR36 exhibited significantly (p = 0.0334) lower amount of GFP-Staufen proteins in the nucleus compared to the controls (Fig 3C). These results suggest that increased accumulation of Staufen in the nucleus by PR toxicity is RNA-dependent.

**Fig. 3.**
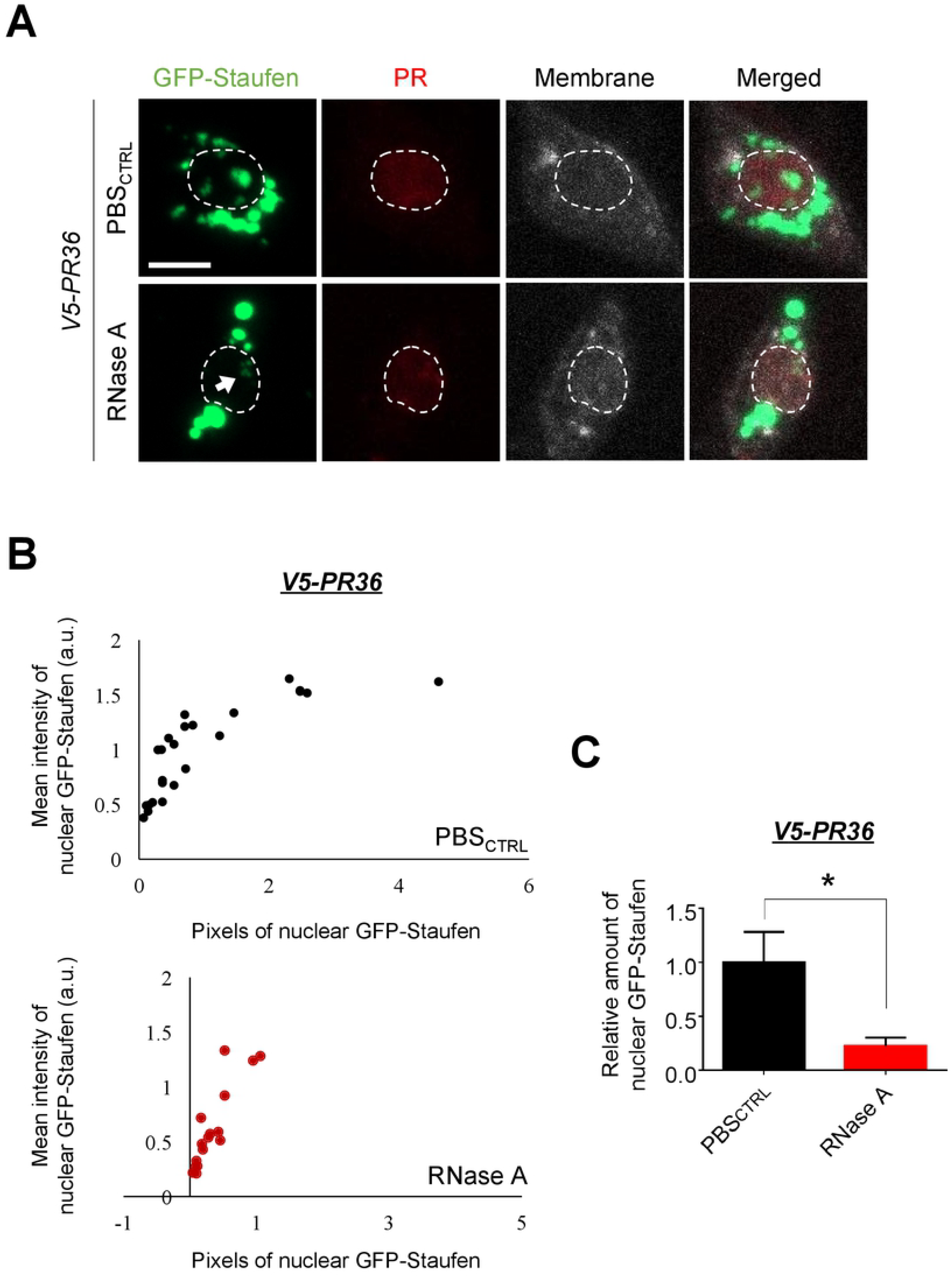
PR toxicity in C4 da neurons induce RNA-dependent accumulation of Staufen in the nucleus. (A) RNase A treatment decreased nuclear accumulation of GFP-Staufen induced by PR toxicity in C4 da neurons (Genotypes. *V5-PR36*: *UAS-GFP-stau/+*; *PPK*^*1a*^-*gal4>UAS-mCD8RFP/UAS-V5-PR36*). Dissected third instar larvae were treated for 1 hr at 37 ℃ with (RNase A) or without (PBS_CTRL_) 20 µg/ml of RNase A. Dashed lines (white) indicate the outlines of the nuclei. Arrow (white) indicates faint nuclear GFP-Staufen signals remained after RNase A treatment. Scale bar, 5 µm. (B) Nuclear GFP-Staufen puncta in C4 da neurons with denoted genotypes from Fig 3A were counted and plotted. y-axis: relative mean intensity of GFP-Staufen puncta in nuclei; x-axis: relative pixel number of GFP-Staufen puncta in nuclei. n≥16 GFP-Staufen puncta. (C) The pixel number and the mean intensity of nuclear GFP-Staufen puncta (obtained from Fig 3B) were multiplied to provide an estimate of relative amount of nuclear GFP-Staufen. *p = 0.0334 by two-tailed student’s t-test; error bars, ± SEM; n≥16 GFP-Staufen puncta.

### Increased nucleolar expression of Fibrillarin by reduced *stau* dosage is a compensatory response to PR toxicity in *Drosophila*

A large portion of heterochromatin, which consists of inactive ribosomal DNA (rDNA), decorates the outer region of the nucleolus within which actively transcribed rDNA are situated [25]. In addition, previous studies have shown that Staufen can localize to the nucleolus [26–28], the dysfunction of which has been reported in C9-ALS/FTD models and patients [29]. These data suggest a possibility that in addition to the heterochromatin, PR toxicity may also induce Staufen to accumulate in or influence the activity of the nucleolus. To test this possibility, we expressed V5-PR36 and GFP-Staufen in C4 da neurons and immunostained for Fibrillarin, which is localized in the dense fibrillar component of the nucleolus [30]. GFP-Staufen and Fibrillarin overlapped only in the fringes (S4A and S4B Fig), but interestingly, we noticed that Fibrillarin staining was considerably stronger (p = 0.0195) in C4 da neurons expressing V5-PR36 than in controls (S5A and S5B Fig). This is consistent with a previous report showing increased nucleolar size determined by Fibrillarin staining in U2OS and NSC-34 cells expressing PR20 [31].

Although Staufen and Fibrillarin overlapped only in the fringes, abnormal accumulation of Staufen in the nucleus may still contribute to the increased staining of Fibrillarin in C4 da neurons expressing V5-PR36. To test this possibility, we examined the amount of Fibrillarin upon V5-PR36 expression in flies with heterozygous mutation (*stau*^*ry9*/+^) for the gene encoding Staufen (*stau*) and compared it to the controls. Surprisingly, heterozygous loss of *stau* actually increased (p = 0.0371) nucleolar staining of Fibrillarin in C4 da neurons expressing V5-PR36 (Fig 4A and 4B). Of note, about 15.6% (5/32) of those neurons showed more than 3.5-fold increase in nucleolar staining of Fibrillarin compared to V5-PR36 only controls. Increased level of Fibrillarin in the nucleolus may be either a toxic feature of or a compensatory response to PR toxicity. To test which is the case, we decreased the level of Fibrillarin by knockdown of the gene encoding Fibrillarin (*Fib*) in *Drosophila* eyes using *GMR*-*gal4* and examined whether PR36-induced retinal degeneration was exacerbated or mitigated. Knockdown of *Fib* exacerbated the PR36-induced retinal degeneration, whereas the knockdown by itself showed no toxicity (Fig 4C). Interestingly, a previous study showed that knockdown of *Fib* can also exacerbate GR-induced retinal degeneration in *Drosophila* [32]. These data suggest that a partial loss of *stau* increases the level of Fibrillarin, which may be a compensatory response to, and not a toxic feature of, PR toxicity.

**Fig. 4.**
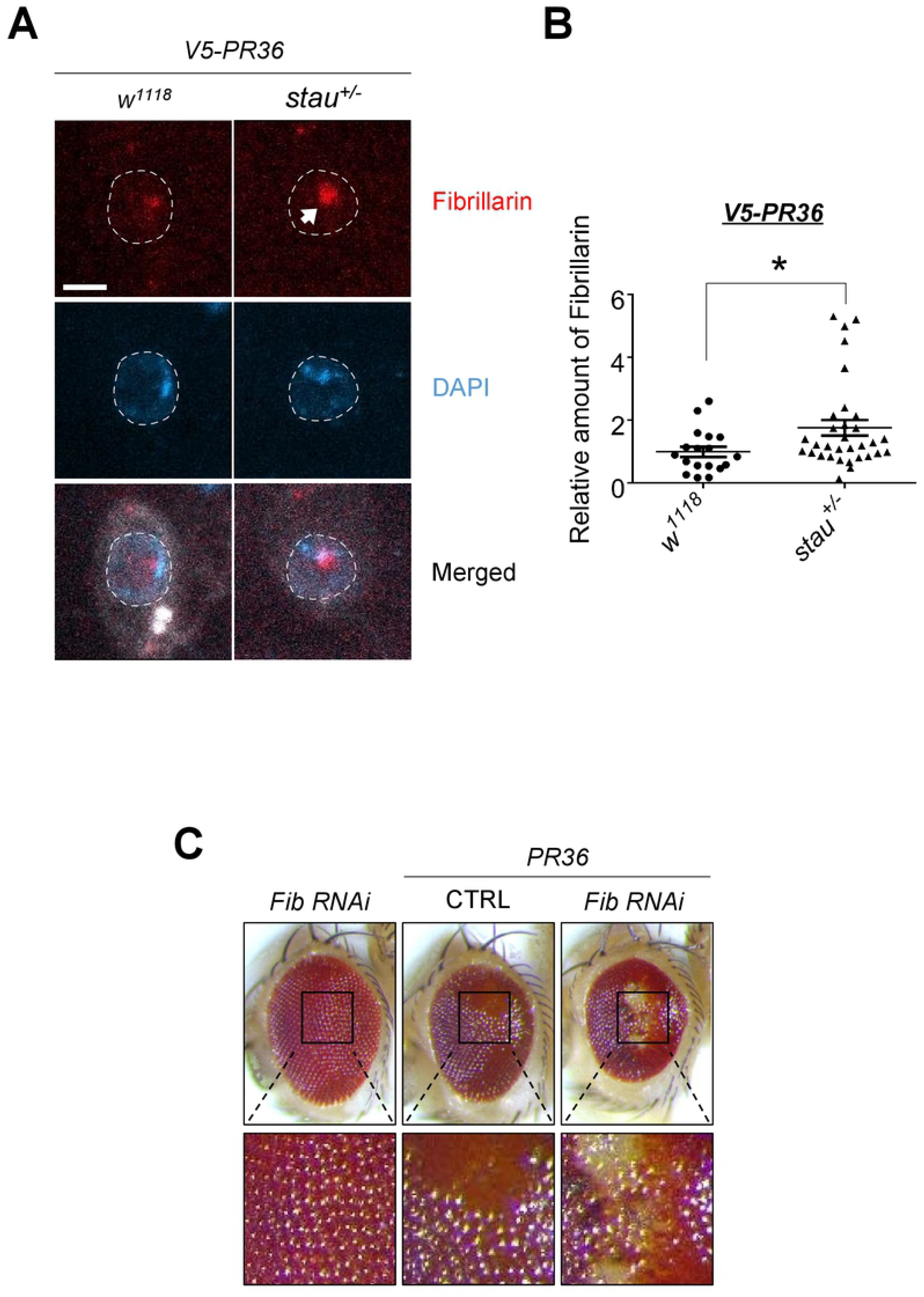
Increased Fibrillarin expression by reduced *stau* dosage is a compensatory response to PR toxicity in *Drosophila*. (A) IHC of dissected third instar larvae shows that heterozygous loss of *stau* increased nucleolar expression of Fibrillarin in V5-PR36-expressing C4 da neurons (Genotypes. *w*^1118^ + *V5-PR36*: *+/+*; *PPK*^*1a*^-*gal4>UAS-mCD8RFP/UAS-V5-PR36*, *stau*^+/−^ + *V5-PR36*: *stau*^*ry9*/+^; *PPK*^*1a*^-*gal4>UAS-mCD8RFP/UAS-V5-PR36*). Dashed lines (white) indicate the outlines of the nuclei. Arrow (white) indicates increased Fibrillarin signal after reducing *stau* dosage. Scale bar, 5 µm. (B) The relative pixel number was multiplied by the relative mean intensity of Fibrillarin puncta to estimate the relative amount of Fibrillarin in C4 da neurons with the denoted genotypes from Fig 4A. *p = 0.0371 by two-tailed student’s t-test; error bars, ± SEM; n≥18 Fibrillarin puncta. (C) Knockdown of *Fib* exacerbated retinal degeneration induced by PR36 toxicity (Genotypes. *Fib RNAi*: *GMR-gal4/UAS-Fib RNAi*, CTRL + *PR36*: *GMR-gal4>UAS-PR36/UAS-40D*, *Fib RNAi* + *PR36*: *GMR-gal4>UAS-PR36/UAS-Fib RNAi*). Fly eyes were imaged at 4 days after eclosion. Images were acquired using 160X objective lens. n=10.

### Heterozygous loss of *stau* rescues PR toxicity in *Drosophila*

We showed above that in neurons expressing V5-PR36, Staufen partially overlaps with Fibrillarin in the nucleolus and that reducing the dosage of *stau* increases the amount of Fibrillarin, which may be a compensatory response to PR toxicity. If this is the case, we hypothesized that reducing *stau* gene dosage might mitigate PR toxicity. To test this hypothesis, we expressed PR36 in the *Drosophila* eyes via *GMR*-*gal4* and compared their external morphologies to the controls. By four days after eclosion, flies expressing PR36 displayed a considerable retinal degeneration with loss of pigments and gain of necrotic spots (Fig 5A). Consistent with our hypothesis, *stau* heterozygous mutation was able to suppress PR36-induced retinal degeneration (Fig 5A).

**Fig. 5.**
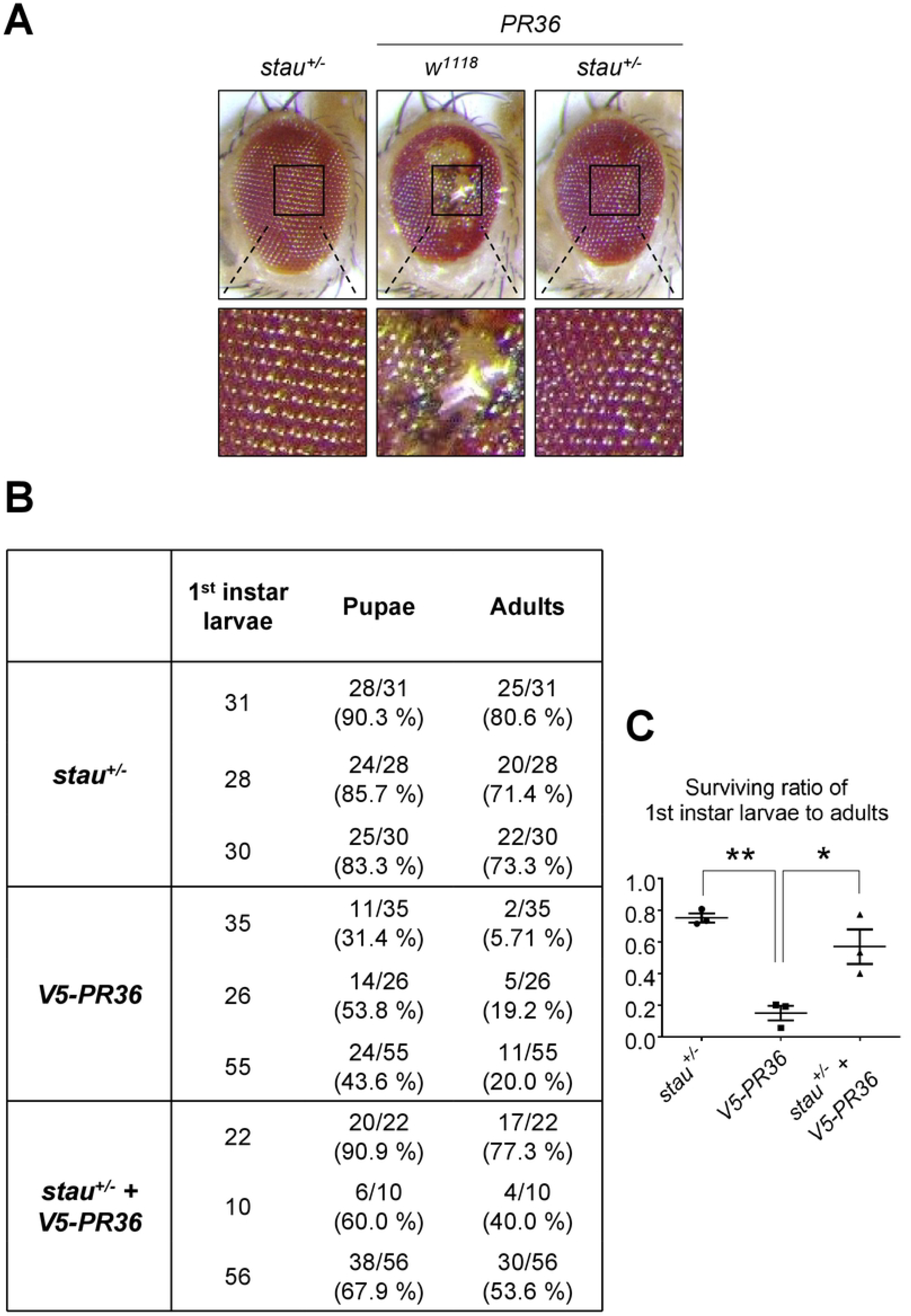
Reduction of *stau* dosage suppresses PR toxicity in *Drosophila*. (A) Heterozygous loss of *stau* restored PR36-induced retinal degeneration (Genotypes. *stau*^*+/−*^: *GMR-gal4/stau*^*ry9*/+^; *+/+*, *w*^1118^ + *PR36*: *GMR-gal4>UAS-PR36*; *+/+*, *stau*^*+/−*^ *+ PR36*: *GMR-gal4>UAS-PR36/stau*^*ry9*/+^; *+/+*). Fly eyes were imaged at 4 days after eclosion. Images were acquired using 160X objective lens. n=9. (B) Heterozygous loss of *stau* restored viability of flies expressing V5-PR36 (Genotypes. *stau*^*+/−*^: *stau*^*ry9*/+^; *elav-gal4/+*, *V5-PR36*: *+/+*; *elav-gal4/UAS-V5-PR36*, *stau*^*+/−*^ + *V5-PR36*: *stau*^*ry9*/+^; *elav-gal4/UAS-V5-PR36*). First instar larvae were collected separately per each line at 25 ℃. Surviving pupae and eclosed adult flies were tallied and listed. Percentages of first instar larvae-to-pupae and first instar larvae-to-adults are denoted in parentheses. (C) The percentages of first instar larva-to-adult viability obtained from three independent experiments were plotted and compared: *stau*^*+/−*^ to V5*-PR36* and *V5-PR36* to *stau*^*+/−*^ + *V5-PR36*. **p = 0.0022; *p = 0.0131 by one-way ANOVA; error bars, ± SEM; n=3 vials.

Next, we asked whether *stau* heterozygous mutation could increase viability of flies expressing V5-PR36. We measured the percentage of larvae (Genotypes. *stau*^*+/−*^: *stau*^*ry9*/+^; *elav-gal4/+*, *V5-PR36*: *+/+*; *elav-gal4/UAS-V5-PR36*, *stau*^*+/−*^ + *V5-PR36*: *stau*^*ry9*/+^; *elav-gal4/UAS-V5-PR36*) reaching the pupal and adult stages and compared them with one another. Most of the *stau*^*+/−*^ larvae reached the pupal (86.46%) and adult (75.14%) stages, whereas only 42.97% and 14.98% of *V5-PR36* larvae reached the pupal and adult stages, respectively (p = 0.0022) (Fig 5B and 5C). Remarkably, the larva-to-pupa (72.92%) and larva-to-adult (56.95%; p = 0.0131) viabilities were largely rescued in *stau*^*+/−*^ + *V5-PR36* larvae (Fig 5B and 5C). Taken together, these results suggest that PR toxicity induces aberrant accumulation of Staufen in the nucleus and that reducing the *stau* gene dosage is sufficient to rescue PR toxicity in *Drosophila*.

## Discussion

Most ALS/FTD pathology can be defined by nuclear loss and cytoplasmic accumulation of disease-associated proteins, such as TDP-43, FUS, TAF15, EWSR1, hnRNPA1, or hnRNPA2/B1, thereby leading to their loss of nuclear function and gain of toxic cytoplasmic function [33]. On the other hand, to our knowledge, no cytoplasmic protein has been identified thus far in ALS/FTD that translocate into the nucleus and accumulate therein. In this study, we identified Staufen, a cytoplasmic dsRBP, as a potential disease-associated protein in C9-ALS/FTD *Drosophila* model that exhibits increased accumulation in the nucleus. We found that, inside the nucleus, Staufen partially interacted with heterochromatin and nucleolus in an RNA-dependent manner. Reducing the dosage of *stau* mitigated retinal degeneration in flies expressing V5-PR36 and rescued their viability.

Among the RBPs screened in this study, only Staufen was identified as a dsRBP. This is a significant point, because what most distinguishes Staufen from other RBPs screened is its ability to bind to dsRNA [34], a toxic byproduct of disrupted heterochromatin caused by PR toxicity [14] and nuclear loss of TDP-43 [13, 23, 24]. In this study, we showed that V5-PR36 expression led to increased co-localization of Staufen with heterochromatin and that degrading RNAs via RNase led to decreased nuclear accumulation of Staufen. These data support the notion that in neurons with V5-PR36 expression, Staufen accumulates around heterochromatin and binds to dsRNA the degradation of which disperses nuclear-accumulated Staufen, leading to its nuclear egress. Nevertheless, whether other dsRBPs also localize near the heterochromatin upon induction of PR toxicity remains to be examined.

In this study, we found that PR expression led to increased staining of Fibrillarin. What might best account for the increased amount of Fibrillarin by PR toxicity? A previous study identified p53 as a repressor of *FBL* (which codes Fibrillarin protein) promoter, thereby regulating the amount of Fibrillarin, in various cancer cell lines [35]. Interestingly, another study showed that dsRNA induces reduction of p53 in HT1080 cells [36]. These pieces of data suggest that increased dsRNA production by PR toxicity may reduce p53, thereby disinhibiting Fibrillarin production. Furthermore, we found that heterozygous mutation of *stau* enhanced the level of Fibrillarin in C4 da neurons expressing V5-PR36, which appears to be a compensatory mechanism against PR toxicity. However, how heterozygous mutation of *stau* induces increased level of Fibrillarin remains unknown. Interestingly, a previous study showed that knockdown of *STAU1* led to reduced half-life of *TP53* (a human homolog of *p53*) mRNA in H1299 cells upon actinomycin D treatment [37]. Based on this study, we speculate that in C4 da neurons expressing V5-PR36, heterozygous *stau* mutation may have decreased p53 level, with subsequent increased Fibrillarin production. In contrast to our speculation, a previous study showed that treatment of a high dose (10μmol/L) of PR_20_ in SH-SY5Y cells led to increased level of p53, albeit with a significant amount of cell death [38]. Thus, further studies are required for a potential role of p53 in mediating Staufen-dependent PR toxicity.

In many neurodegenerative diseases, such as Huntington’s disease, abnormal nuclear accumulation of proteins is known to be highly toxic [39, 40]. Although whether nuclear-accumulated Staufen is toxic in C9-ALS/FTD remains disputable, our results nevertheless are largely consistent with the notion that nuclear-accumulated Staufen can be toxic. Out results show that 1) Staufen accumulates in the nucleus and encroaches on the heterochromatin and nucleolus, possibly affecting their functions, 2) reducing *stau* dosage in PR-expressing neurons increased the amount of Fibrillarin, which may be a compensatory response to PR toxicity, and 3) *stau* heterozygous mutation rescues retinal degeneration and viability. In addition, a recent proteomics study identified Stau2, an ortholog of *Drosophila* Staufen [41], to be commonly enriched in both PR and GR interactome in rat primary neurons [42]. Consistent with the previous study, we found that V5-PR36, which localizes predominantly to the nucleus, interacts with Staufen in *Drosophila* neurons (S6A Fig). This raises the possibility that Staufen may interact with PR in the nucleus in an RNA-dependent manner to contribute to toxicity. Finally, Staufen in the form of puncta in the nucleus may sequester various proteins and RNAs whose functions may be disrupted. On the other hand, Staufen puncta in the nucleus nearby heterochromatin and nucleolus may be benign; and reduction of *stau* may have rescued retinal degeneration and viability through reducing the level of cytoplasmic Staufen. Therefore, further in-depth study is necessary to clearly differentiate between nuclear and cytoplasmic Staufen-mediated toxicity.

A recent study showed that Spinocerebellar ataxia type 2 (SCA2) mouse model and human patients presented elevated level of Stau1 protein whose interaction with polyQ-expanded ATXN2 caused neurotoxicity [43]. Reducing *STAU1* in SCA2 mouse improved motor behaviors and reduced polyQ-expanded ATXN2 aggregation. In the same study, they also showed an elevated level of Stau1 in fibroblasts from ALS patients with TDP-43 G298S mutation [43]. However, whether increased Stau1 level in the fibroblasts of ALS patients are toxic has not been explored. In our study, we showed that Staufen accumulates in the nucleus and that reducing *stau* suppressed retinal degeneration and rescued viability of flies exhibiting PR toxicity. Taken together, these data suggest that aberrant accumulation of Staufen in neurons is toxic and that its reduction might be a suitable therapeutic target for a spectrum of diseases including SCA2, ALS, and FTD.

## Materials and methods

### *Drosophila* stocks

All flies were maintained at 25 ℃ and 60 % humidity. The following lines were obtained from Bloomington *Drosophila* Stock Center (Indiana, USA): *w*^1118^(3605); *UAS-G_4_C_2_36* (58688); *UAS-G_4_C_2_36-RNA only* (58689); *UAS-PR36* (58694); *RFP-HP1* (30562); *stau*^*ry9/+*^ (10742); *UAS-TBPH RNAi* (29517); *UAS-Luciferase* (35788); *elav-GeneSwitch(GS)-gal4* (43642); *D42-gal4* (8816); *GMR-gal4* (1104); and *elav-gal4* (8760). The following lines were obtained from Vienna *Drosophila* Resource Center (VDRC): *UAS-Fib RNAi* (104372KK) and *UAS-40D* (60101). *UAS-GFP-stau* was provided by Andrea Brand (University of Cambridge, UK); *UAS-mCD8RFP* and *PPK*^*1a*^-*gal4* were provided by Yuh Nung Jan (UCSF, USA). *ElavGS-gal4* expression was induced by feeding adult flies 100 μM of RU486 (Mifepristone) (Sigma. M8046) for 20 days after eclosion at 27 ℃.

### Molecular cloning and generation of transgenic flies

V5-PR36, HA-GR36, and Myc-PA36 were subcloned (Genscript, Inc.) into pACU2 vectors (from Chun Han, Cornell University) using BglIII and XhoI sites, and the epitopes were added at the N-terminus. Transgenic flies were generated by BestGene, Inc. The nucleotide sequences of the generated constructs are described below:

#### *BglII*-V5-PR36-*XhoI*

5’-*CACAGATCTCCACC***ATGGGTAAGCCTATCCCTAACCCTCTCCTCGGTCTCGATT CTACGGGTGGAGGCGGTAGTCCTCGACCTAGACCAAGACCTCGGCCTAGGC CACGTCCTAGACCACGTCCTAGGCCACGACCTCGTCCTCGACCACGTCCTC GTCCTAGGCCCAGACCACGCCCTAGACCAAGACCCAGACCACGACCACGGC CTCGACCTAGGCCTCGTCCACGACCTCGCCCAAGACCTCGTCCGAGACCTA GGCCCAGACCTCGTCCTAGACCAAGACCCAGA***TGATAACTCGAGGCA*-3’

#### *BglII*-HA-GR36-*XhoI*

5’-*CACAGATCTCCACC***ATGTACCCATACGATGTTCCAGATTACGCTGGTGGAGGC GGTAGTGGTCGAGGACGTGGTCGAGGAAGAGGTCGTGGTCGTGGACGAGG TAGAGGACGTGGAAGAGGTCGAGGACGTGGTAGAGGTCGAGGAAGAGGTC GTGGACGAGGACGTGGTCGAGGAAGAGGTCGTGGTCGTGGACGAGGTAGA GGACGTGGAAGAGGTCGAGGACGTGGTAGAGGTCGAGGAAGAGGTCGTGG ACGAGGACGTGGTCGAGGAAGA***TGATAACTCGAGGCA*-3’

#### *BglII*-Myc-PA36-*XhoI*

5’-*CACAGATCTCCACC***ATGGAACAAAAACTCATCTCAGAAGAGGATCTGGGTGGA GGCGGTAGTCCTGCCCCTGCTCCAGCTCCAGCACCTGCACCAGCCCCAGCA CCTGCTCCAGCACCCGCGCCTGCACCAGCTCCAGCACCGGCACCTGCTCCG GCACCTGCACCTGCGCCAGCTCCTGCTCCTGCGCCAGCTCCAGCACCAGCA CCTGCTCCAGCTCCAGCTCCTGCACCAGCACCCGCACCTGCTCCAGCCCCA GCTCCTGCACCTGCTCCTGCA***TGATAACTCGAGGCA*-3’

### Immunohistochemistry (IHC)

Third instar larvae were dissected and immediately fixed in 3.7 % formaldehyde (JUNSEI, #50-00-0) for 20 min at room temperature (RT). Formaldehyde was diluted in 1X phosphate buffered saline (PBS) (MENTOS). After washing three times at RT in 0.3 % PBST (0.3% triton X-100 in 1X PBS), the samples were incubated in the 5 % normal donkey serum (NDS) diluted in washing buffer (blocking buffer) for 1 hr at RT. Next, the samples were incubated in primary antibodies diluted in blocking buffer overnight at 4 ℃. To detect GFP-Staufen, V5-PR36, and endogenous Fibrillarin, the following primary antibodies were used: rabbit anti-GFP (Abcam, ab183734) (1:500); mouse anti-V5 (Thermo Scientific, R960-25) (1:500); and mouse anti-Fibrillarin (Abcam, ab4566) (1:200). After washing five times for 10 min in washing buffer at RT, the samples were incubated with secondary antibodies diluted in blocking buffer for 2-4 hours at RT. The following secondary antibodies were used: goat anti-rabbit Alexa 488 (Invitrogen, A11034) (1:1000); goat anti-mouse Alexa 647 (Invitrogen, A21236) (1:1000 or 1:400); and goat anti-HRP Cy3 (Jackson Immunoresearch Laboratories, #123-165-021) (1:200). The anti-HRP antibody was used to label the whole neuronal membranes of soma and dendrites. The samples were then washed five times for 5 min in washing buffer and mounted using mounting medium with DAPI (4’,6-diamidino-2-phenylindole) (VECTASHIELD, H-1200) for imaging. DAPI was used for visualizing heterochromatin located in the nucleus. For IHC of adult brains and larval brains, all primary antibodies were diluted 1:200 and all secondary antibodies were diluted 1:400 in blocking buffer. All procedures in IHC experiments were performed in the dark.

### Imaging analysis

Confocal images were acquired using LSM 780 (ZEISS) or LSM 800 (ZEISS) microscope. *In vivo* neuronal images of 3^rd^ instar larvae were taken at 200x magnification with the confocal microscope. Neuronal images from staining experiments in 3^rd^ instar larvae were taken at 400x magnification with the confocal microscope. All C4 da neurons were acquired from the abdominal segments A4-A6. Retinal images were acquired using Leica SP5. Fly eyes (left eyes only) were taken at 160x magnification immediately upon dissection.

### Ribonuclease A (RNase A) treatment

Samples were incubated in 20 μg/ml of RNase A (Sigma, R6513) suspended in 1X PBS for 1 hr at 37 ℃ before fixation. As controls, samples were incubated in 1X PBS instead of RNase A for 1 hr at 37 ℃ before fixation. The next steps of IHC were performed as described above.

### RNA Fluorescent *in situ* hybridization (FISH)

The protocol for protein-RNA double labeling in *Drosophila* ovaries [44] was slightly modified to apply FISH to C4 da neurons of third instar larval fillet. To confirm mRNA reduction in RNase A-treated C4 da neurons of third instar larvae, a Cy 5-tagged poly(A)-tail-targeting (Cy5-oligo-dT) probe was designed as follows: 5’-Cy5-TTT TTT TTT TTT TTT TTT TT-3’ (6555.6 g/mol MW, 37.9 ℃ Tm) (Macrogen Inc.). All procedures in FISH experiments were performed in the dark. In FISH experiments, IHC was first performed using dissected third instar larvae, but with different solutions from IHC described above: 0.1 % tween-20 (Biotech, #9005-64-5) diluted in 1X PBS (PBT); 3.7 % formaldehyde in PBT; and 1 % skim milk in PBT. Then, FISH procedures were applied to the samples as previously described [44]. 3 μM (tests for 0.1-5 μM) of a Cy5-oligo-dT probe was hybridized to the samples at 37.9 ℃ (Tm) overnight. The following FISH solutions were used: EtOH (Merk, #64-17-5); Xylenes (Sigma, #1330-20-7); RIPA (Biosesang, R2002); Formamide (Sigma, F9037); 5X SSC (Biosesang, S2012); Heparin (Sigma, H4784); DEPC-treated water (MENTOS, M1409). Finally, the samples were mounted for imaging.

### Co-immunoprecipitation (co-IP)

Co-IP experiments were performed as previously described [45] with some modifications listed below. More than 250 fly heads were collected from each line and suspended in 400 μl lysis buffer (25mM Tris-buffered saline (Tris-HCl) pH 7.5, 150mM NaCl, 0.1 % tween-20, 1mM EDTA, 10 % Glycerol) with 1:100 ratio of protease inhibitor cocktail (Thermo Scientific, #87786). Samples were completely homogenized and then centrifuged at 13,300 rpm for 20 min at 4 ℃. Each sample containing equal amount of proteins was incubated overnight at 4 ℃ with mouse anti-V5 (1:500; Thermo Scientific, R960-25). Then, protein A/G plus-agarose beads (Santa Cruz, sc-2003) were added, and samples were further incubated for 3 hr at 4 ℃ on a nutator. After the samples were centrifuged three times at 3,200 rpm for 5 sec at 4 ℃, the supernatant was discarded. The primary and secondary antibodies used to detect the bands are as follows: mouse anti-GFP (Santa Cruz, sc-9996) (1:1000); mouse anti-V5 (Thermo Scientific, R960-25) (1:1000); mouse anti-beta-tubulin (DSHB, E7-s) (1:1000); and goat anti-mouse IgG-HRP (Santa Cruz, sc-3697) (1:2000).

### Image processing and statistical analysis

Obtained confocal images were further analyzed using Image J for quantification of pixel number and mean intensity. Microsoft Excel was used to plot puncta patterns on the graph. For statistical analyses, GraphPad Prism 6.01 was used for two-tailed student’s t-test and one-way ANOVA. Relative mean intensities and pixel values were calculated and used for analyses. Error bars show SEM and all p values were summarized with asterisks: *p < 0.05; **p < 0.05; ***p < 0.005; ****p < 0.0005.

## Acknowledgements and Funding

This work was funded by the Ministry of Science, Information and Communications Technology (ICT) & Future Planning (19-BR-03-03) (to CMH), the KBRI Research Program of the Ministry of Science, ICT & Future Planning (19-BR-02-04) (to JK) (19-BR-02-03) (to H-JK), and Basic Science Research Program through the National Research Foundation of Korea, funded by the Ministry of Science and ICT (2018R1A2B6001607) (to SBL); the Development of Platform Technology for Innovative Medical Measurements Program from the Korea Research Institute of Standards and Science Grant (KRISS-2019-GP2019-0018) (to SBL). The funders had no role in study design, data collection and analysis, decision to publish, or preparation of the manuscript.

## Author Contributions

### Conceptualization

Sung Bae Lee, Eun Seon Kim, Chang Geon Chung

### Data curation

Eun Seon Kim, Chang Geon Chung, Yoon Ha Kim

### Formal analysis

Eun Seon Kim, Chang Geon Chung, Yoon Ha Kim, In Jun Cha, Jeong Hyang Park, Hyung-Jun Kim, Chang Man Han, Jaekwang Kim

### Funding acquisition

Sung Bae Lee, Hyung-Jun Kim, Chang Man Han, Jaekwang Kim

### Investigation

Eun Seon Kim, Chang Geon Chung, Yoon Ha Kim

### Methodology

Eun Seon Kim, Chang Geon Chung, Yoon Ha Kim, In Jun Cha, Jeong Hyang Park

### Project Administration

Sung Bae Lee

### Resources

Sung Bae Lee

### Software

Eun Seon Kim, Chang Geon Chung, Yoon Ha Kim

### Supervision

Sung Bae Lee

### Validation

Eun Seon Kim, Chang Geon Chung, Yoon Ha Kim

### Visualization

Eun Seon Kim, Chang Geon Chung, Yoon Ha Kim

### Writing—Original Draft Preparation

Eun Seon Kim, Chang Geon Chung

### Writing—Review & Editing

Sung Bae Lee, Eun Seon Kim, Chang Geon Chung

## Supporting information

**S1 Fig. Arginine-rich DPR-expressing constructs increase nuclear accumulation of Staufen.**

(A) *In vivo* imaging of *Drosophila* third instar larval C4 da neurons shows that *C9orf72* constructs expressing arginine-rich DPRs increased nuclear accumulation of GFP-Staufen (Genotypes. CTRL: *UAS-GFP-stau/+*; *PPK*^*1a*^-*gal4>UAS-mCD8RFP/+*, *G_4_C_2_36*: *UAS-GFP-stau/UAS-G_4_C_2_36*; *PPK*^*1a*^-*gal4>UAS-mCD8RFP/+*, *G_4_C_2_36-RNA only*: *UAS-GFP-stau/UAS-G_4_C_2_36-RNA only*; *PPK*^*1a*^-*gal4>UAS-mCD8RFP/+*, *Myc-PA36*: *UAS-GFP-stau/+*; *PPK*^*1a*^-*gal4>UAS-mCD8RFP/UAS-Myc-PA36*, *HA-GR36*: *UAS-GFP-stau/+*; *PPK*^*1a*^-*gal4>UAS-mCD8RFP/UAS-HA-GR36*). Magnified images of the nuclei of left panels are presented at the bottom of each image; scale bar (yellow), 2 µm. Dashed circular lines (white) outline the nuclei. Arrows (white) indicate stronger nuclear GFP-Staufen signals compared to the controls. Scale bar (white), 10 µm. (B) Nuclear GFP-Staufen puncta in C4 da neurons with denoted genotypes from S1A Fig were counted and plotted. y-axis: relative mean intensity of GFP-Staufen puncta in nucleus; x-axis: relative pixel number of GFP-Staufen puncta in nucleus. n≥ 17 GFP-Staufen puncta. (C) The pixel number and the mean intensity of nuclear GFP-Staufen puncta (obtained from S1B Fig) were multiplied to provide an estimate of relative amount of nuclear GFP-Staufen. ns: not significant; **p = 0.0037; ***p = 0.0004 by two-tailed student’s t-test; error bars, ± SEM; n ≥17 GFP-Staufen puncta.

**S2 Fig. *TBPH* knockdown increases nuclear accumulation of Staufen in neurons.**

(A) *In vivo* imaging shows that *TBPH* RNAi increases nuclear accumulation of Staufen in C4 da neurons of third instar larvae (Genotypes. CTRL: *UAS-GFP-stau/+*; *PPK*^*1a*^-*gal4>UAS-mCD8RFP/UAS-Luciferase*, *TBPH RNAi*: *UAS-GFP-stau/+*; *PPK*^*1a*^-*gal4>UAS-mCD8RFP/UAS-TBPH RNAi*). Magnified images of the nuclei in the left panels are presented at the bottom of images; scale bar (yellow), 1 µm. Dashed circular lines (white) outline the nuclei. Arrow (white) indicates stronger nuclear GFP-Staufen signal compared to the control. Scale bar (white), 10 µm; (yellow), 1 µm. (B) Nuclear GFP-Staufen puncta in C4 da neurons with denoted genotypes from S2A Fig were counted and plotted. y-axis: relative mean intensity of GFP-Staufen puncta in nucleus; x-axis: relative pixel number of GFP-Staufen puncta in nucleus. n≥57 GFP-Staufen puncta. (C) The pixel number and the mean intensity of nuclear GFP-Staufen puncta (obtained from S2B Fig) were multiplied to provide an estimate of relative amount of nuclear GFP-Staufen. ****p < 0.0001 by two-tailed student’s t-test; error bars, ± SEM; n≥57 GFP-Staufen puncta. (D) The numbers of nuclear GFP-Staufen puncta (obtained from S2B Fig) were counted and compared. *p = 0.0179 by two-tailed student’s t-test; error bars, ± SEM; n=10 neurons.

**S3 Fig. RNase A treatment efficiently reduces RNA levels detected via RNA FISH.**

(A) RNA FISH of dissected third instar larvae confirms the effect of RNase A treatment on decreasing mRNA amount in V5-PR36-expressing C4 da neurons (Genotype. *V5-PR36*: *UAS-GFP-stau/+*; *PPK*^*1a*^-*gal4>UAS-mCD8RFP/UAS-V5-PR36*). 3 µM of Cy5-oligo-dT DNA probe was used to detect mRNA. Dissected third instar larvae were treated for 1 hr at 37 ℃ with (RNase A) or without (PBS_CTRL_) 20 µg/ml of RNase A. Dashed circular lines (white) outline the nuclei. Scale bar (white), 5 µm. (B) Intensity profiles of fluorescent signals representing nuclear GFP-Staufen (Green) and mRNA (Red) show decreased nuclear accumulation of GFP-Staufen after RNase A treatment. y-axis: intensity of fluorescent signals; x-axis: distance (µm). Top right insets are the images used for measuring intensity profiles of fluorescent signals; solid line (yellow) is x-axis (distance). Scale bar (white), 2 µm.

**S4 Fig. Nuclear-accumulated Staufen is contiguous to Fibrillarin that is increased by PR toxicity in C4 da neurons.**

(A) IHC of dissected third instar larvae shows that nuclear-accumulated GFP-Staufen is contiguous to Fibrillarin, the expression of which was increased in V5-PR36-expressing C4 da neurons (Genotypes. CTRL: *UAS-GFP-stau/+; PPK*^*1a*^-*gal4>UAS-mCD8RFP/+*, *V5-PR36*: *UAS-GFP-stau/+; PPK*^*1a*^-*gal4>UAS-mCD8RFP/UAS-V5-PR36*). Dashed circular lines (white) outline the nuclei. Arrow (white) indicates conjunction of nuclear-accumulated GFP-Staufen with Fibrillarin. Scale bar, 5 µm. (B) 3D images of S4A Fig are presented to better visualize the conjunction of GFP-Staufen with Fibrillarin in the nucleus. Arrows (white) indicate the overlap between nuclear-accumulated GFP-Staufen (green) and Fibrillarin (white).

**S5 Fig. PR toxicity increases nucleolar expression of Fibrillarin in neurons.**

(A) IHC shows that nucleolar expression of Fibrillarin is increased by V5-PR36 expression in C4 da neurons (Genotypes. CTRL: *+/+; PPK*^*1a*^-*gal4>UAS-mCD8RFP/UAS-pACU2-empty*, *V5-PR36*: *+/+; PPK*^*1a*^-*gal4>UAS-mCD8RFP/UAS-V5-PR36*). Dashed lines (white) indicate the outlines of the nuclei. Arrow (white) indicates stronger Fibrillarin signal by PR toxicity compared to the control. Scale bar, 5 µm. (B) The pixel number and the mean intensity of Fibrillarin (obtained from S5A Fig) were multiplied to provide an estimate of relative amount of Fibrillarin. *p = 0.0195 by two-tailed student’s t-test; error bars, ± SEM; n≥7 Fibrillarin puncta.

**S6 Fig. Staufen physically interacts with PR in *Drosophila* brains.**

(A) Co-IP shows that GFP-Staufen forms a complex with V5-PR36 in V5-PR36-expressing adult fly brains (Genotypes. *V5-PR36*: *+/+*; *elavGS-gal4/UAS-V5-PR36*, *GFP-stau* + *V5-PR36*: *UAS-GFP-stau /+*; *elavGS-gal4/UAS-V5-PR36*). The transgenic expression was induced by feeding 100 µM of RU486 for 20 days after eclosion at 27 ℃. β-tubulin was used as an internal control. n=3.

